# A Cre-dependent reporter mouse for quantitative real-time imaging of Protein Kinase A activity dynamics

**DOI:** 10.1101/2023.10.31.565028

**Authors:** Elizabeth I. Tilden, Aditi Maduskar, Anna Oldenborg, Bernardo L. Sabatini, Yao Chen

**Affiliations:** Department of Neuroscience, Washington University in St. Louis, St. Louis, MO, United States; Ph. D. Program in Neuroscience, Washington University in St. Louis; Howard Hughes Medical Institute, Department of Neurobiology, Harvard Medical School, Boston, MA, United States

## Abstract

Intracellular signaling dynamics play a crucial role in cell function. Protein kinase A (PKA) is a key signaling molecule that has diverse functions, from regulating metabolism and brain activity to guiding development and cancer progression. We previously developed an optical reporter, FLIM-AKAR, that allows for quantitative imaging of PKA activity via fluorescence lifetime imaging microscopy and photometry. However, using viral infection or electroporation for the delivery of FLIM-AKAR is invasive, cannot easily target sparse or hard-to-transfect/infect cell types, and results in variable expression. Here, we developed a reporter mouse, *FL-AK*, which expresses FLIM-AKAR in a *Cre*-dependent manner from the *ROSA26* locus. *FL-AK* provides robust and consistent expression of FLIM-AKAR over time. Functionally, the mouse line reports an increase in PKA activity in response to activation of both G_αs_ and G_αq_-coupled receptors in brain slices. *In vivo, FL-AK* reports PKA phosphorylation in response to neuromodulator receptor activation. Thus, *FL-AK* provides a quantitative, robust, and flexible method to reveal the dynamics of PKA activity in diverse cell types.

## Introduction

Multiple studies in recent years have shown that cells can encode and decode information through the spatial and temporal dynamics of intracellular signals^1^. Transient, sustained, or oscillatory patterns of the same intracellular signal, for example, can result in distinct outcomes including proliferation, differentiation, cell death, or cell cycle arrests^2–8^. The importance of signal dynamics has been demonstrated in cell biology, development, immunology, cancer biology, and neuroscience.

Protein phosphorylation is a widely used signal transduction process and is catalyzed by protein kinases. Protein kinase A (PKA) is a ubiquitous and functionally important protein kinase. In the nervous system, it integrates inputs from many extracellular signals and has profound effects on neuronal excitability, synaptic transmission, synaptic plasticity, and learning and memory^9–26^. In cancer biology, it controls oncogenic signaling and regulates mesenchymal-to-epithelial transition^27,28^. Furthermore, PKA activity is critical in multiple processes in metabolism, development, vascular biology, pancreatic, and kidney functions^29–33^. Moreover, the dynamics of PKA activity are critical for its functions. On a cellular level, distinct PKA regulation by different receptors in different cell types results in cell-type specific and learning stage-specific contributions to learning^13^. On a subcellular level, PKA activation in different compartments or microdomains regulates distinct functions such as differential modulation of receptors and channels^34^. Temporally, different duration of PKA phosphorylation can lead to distinct modes of temporal integration, resulting in control of mating duration^24,35^. Thus, being able to watch PKA activity with cellular resolution in real time is critical to understand PKA function.

Because of the importance of PKA dynamics, multiple optical reporters of PKA activity have been made^36–41^. Together, these sensors have revealed the importance of the PKA dynamics that underlie lipid metabolism, cancer biology, learning, movement, and response to chemicals in the brain that modulate the nervous system (neuromodulators)^10,13,25–27,32,35–38,42^. They are all based on an original ratiometric Förster Resonance Energy Transfer (FRET) reporter developed by Jin Zhang and Roger Tsien^36^, which works successfully in cells but has limitation in thick brain tissue, especially due to the challenge of using FRET with two photon (2p) microscopy. To make the original sensor compatible with 2p imaging in thick brain tissue and *in vivo*, we converted it into one that is compatible with two photon fluorescence lifetime imaging microscopy (2pFLIM). We named it FLIM-compatible A Kinase Activity Reporter (FLIM-AKAR)^39^. FLIM, which measures the time it takes between excitation and emission of light from the donor fluorophore, offers excellent signal to noise ratio and is especially important for compatibility with 2p microscopy^39,43,44^.

Previously, PKA activity sensors were delivered *in vivo* via viral infection or *in utero* electroporation (IUE) of DNA. Although these methods are effective in sensor delivery, they result in expression level variation from cell to cell, and from mouse to mouse. Furthermore, region-specific delivery of sensors via IUE or virus makes it hard to target sparsely distributed cell types. Finally, the surgeries required involve additional experimental time, are invasive, and cause inflammation afterwards. For other sensors, such as the calcium sensor GCaMP, the development of knock-in mouse lines have benefited the scientific community tremendously with consistent expression, ease of targeting to rare cell types, and removal of the need for surgeries^45–47^.

To overcome the limitations of sensor delivery with IUE and virus, we constructed a knock-in reporter mouse of PKA activity that expresses FLIM-AKAR in a *Cre* recombinase-dependent way. We find robust and consistent expression of FLIM-AKAR in multiple cell types in the brain. We demonstrate successful reporting of PKA activity in response to neuromodulator G protein-coupled receptor (GPCR) activation in specific cell types in brain slices and in freely moving mice. Thus, the PKA activity reporter mouse line can be deployed to reveal PKA dynamics in genetically identifiable cell types in the brain and beyond.

## Results

To characterize the utility of the *FLIM-AKAR*^*flox/flox*^ (*FL-AK*) knock-in mouse line, we crossed *FL-AK* mice with *Cre* lines to express FLIM-AKAR in selected cell populations in the brain. We labelled Type 1 dopamine receptor (D1R) expressing spiny projection neurons (D1R-SPNs) by crossing *FL-AK* with *Tg(Drd1a-cre)* mice^48–50^. We labelled excitatory neurons (and a small subset of glia) in the neocortex and hippocampus by crossing *FL-AK* mice with *Emx1*^*IREScre*^ mice^51^. We then imaged FLIM-AKAR responses to neuromodulator receptor activation in brain slices and in freely behaving mice.

### *FLIM-AKAR*^*flox/flox*^ mice show robust and steady expression levels over time

To achieve consistent expression of the PKA activity reporter FLIM-AKAR, we generated a knock-in mouse of *Cre* recombinase-dependent *FLIM-AKAR* in the *ROSA26* locus (Fig. 1a). Here, the *FLIM-AKAR* gene is under the control of the cytomegalovirus early enhancer/chicken β-actin *(CAG)* promoter. The addition of a *lox-stop-lox* cassette makes the reporter gene *Cre*-dependent, allowing for selective expression of FLIM-AKAR in specific cell types. FLIM-AKAR is a fluorescence lifetime-based, genetically encoded optical reporter of PKA activity^39^. It consists of a donor fluorophore of monomeric enhanced green fluorescent protein (meGFP) and an acceptor fluorophore of dark yellow fluorescent protein (sREACH). When PKA phosphorylates the substrate consensus region within the linker region, the resulting phosphopeptide binds to the FHA1 phosphopeptide binding domain that is also in the linker region, thus bringing the donor and acceptor fluorophores closer together. This results in increased FRET and decreased fluorescence lifetime. This conformational change can also be reversed by phosphatases, which release the phosphopeptide from its binding domain through dephosphorylation (Fig. 1b). Thus, FLIM-AKAR is a phosphorylation substrate reporter that reports the balance between PKA and phosphatase. After crossing *FL-AK* mice to *Cre* lines to express FLIM-AKAR in selected cell types, we assessed FLIM-AKAR expression and functional responses with 2pFLIM.

**Figure 1.**
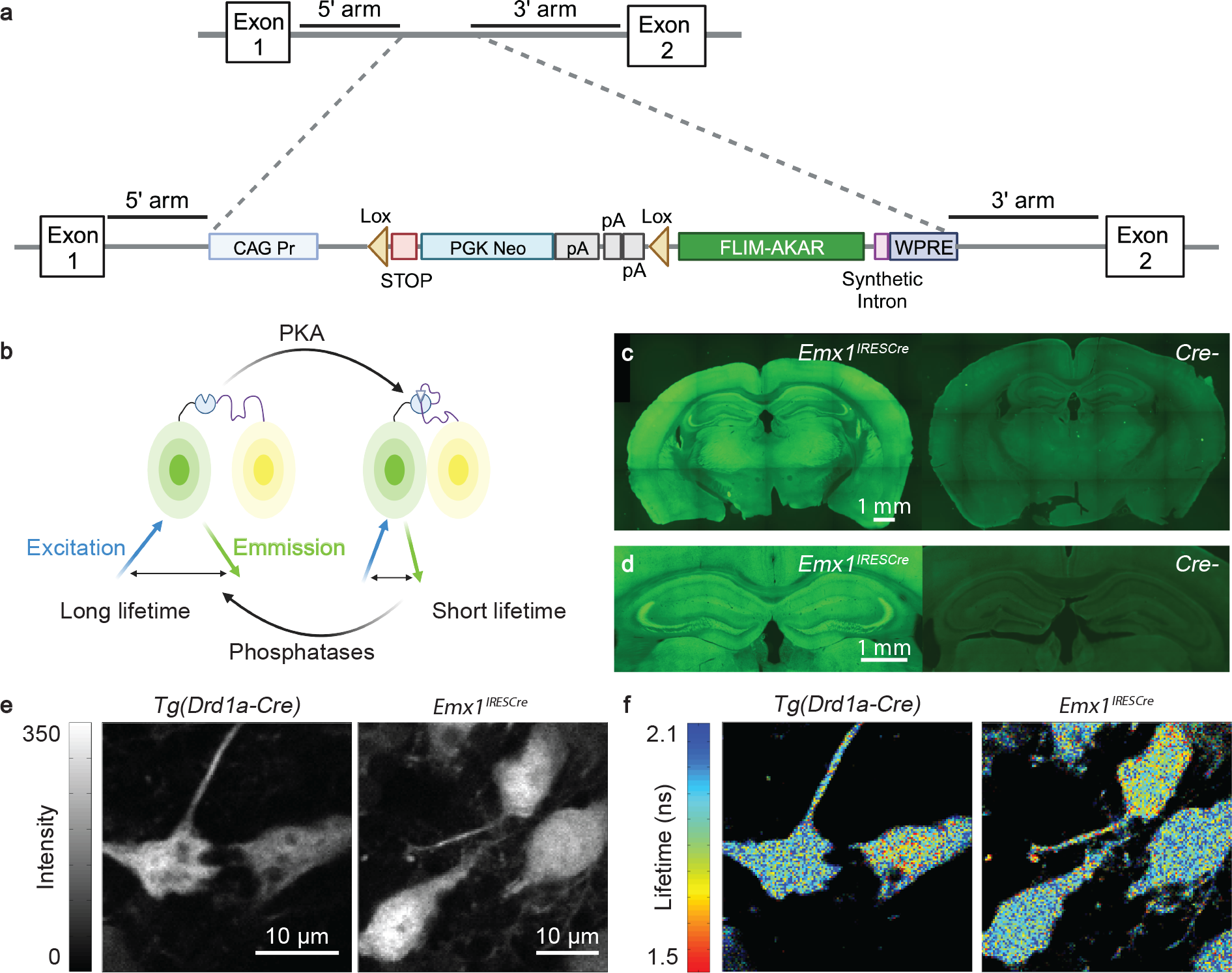
Expression of FLIM-AKAR across multiple cell types in the *FLIM-AKAR*^*flox/flox*^ mouse. **(a)** Schematic of the gene targeting strategy to generate the *FLIM-AKAR*^*flox/flox*^ mice. **(b)** Schematic of how FLIM-AKAR works. FLIM-AKAR detects PKA phosphorylation via the change of FRET between a donor fluorophore and an acceptor fluorophore. When PKA phosphorylates the PKA consensus substrate region in the linker, the two fluorophores come closer together, resulting in increased FRET and decreased fluorescence lifetime. The process can be reversed by phosphatases. **(c-d)** Images of coronal slices showing FLIM-AKAR expression in the whole brain (c) and hippocampus (d) in *Emx1*^*IRESCre*^*;FLIM-AKAR*^*flox/flox*^ (left) and *Cre-;FLIM-AKAR*^*flox/flox*^ (right) mice. Images within the same panel have matching imaging conditions. **(e-f)** 2p fluorescence intensity (e) and lifetime (f) images of D1R-SPNs in the dorsal striatum (left) and pyramidal neurons in CA1 (right).

*Emx1*^*IREScre*^;*FLIM-AKAR*^*flox/flox*^ mice showed robust FLIM-AKAR expression across the cortex and in the hippocampus (Fig. 1c,d). In contrast, in *Cre-/-*; *FLIM-AKAR*^*flox/flox*^ animals, there was very little green signal, demonstrating the *Cre* dependence of FLIM-AKAR expression (Fig. 1c,d). To observe cellular-level FLIM-AKAR expression across cell types, we collected both fluorescence intensity and lifetime data from acute brain slices using 2pFLIM. We observed reliable FLIM-AKAR expression throughout the cell in D1R-SPNs in *FL-AK* reporter mice crossed with the *Tg(Drd1-cre)* line^48–50^ and in CA1 pyramidal neurons of *FL-AK* reporter mice crossed with the *Emx1*^*IREScre*^ line^51^ (Fig. 1e). Interestingly, these neurons also displayed heterogeneity of fluorescence lifetime, indicating different PKA phosphorylation states between subcellular compartments and between cells (Fig. 1f)

In order to determine whether *FL-AK* reporter mice show stable expression of FLIM-AKAR over time, we performed 2p imaging of acute striatal slices from *Tg(Drd1a-Cre); FLIM-AKAR*^*flox/flox*^ mice across a range of ages (Fig. 2). In both the nuclear and cytoplasmic compartments, expression level of FLIM-AKAR was not significantly different between mice aged 0-6 weeks (n = 14 cells from 4 mice) and mice aged 7-14 weeks (n = 16 cells from 6 mice) (Fig. 2c,d), nucleus: p=0.755, cytoplasm: p= 0.787; 2-tailed Mann-Whitney U test). Thus, *FL-AK* reporter mice show robust and consistent expression and can be used to determine PKA activity across ages.

**Figure 2.**
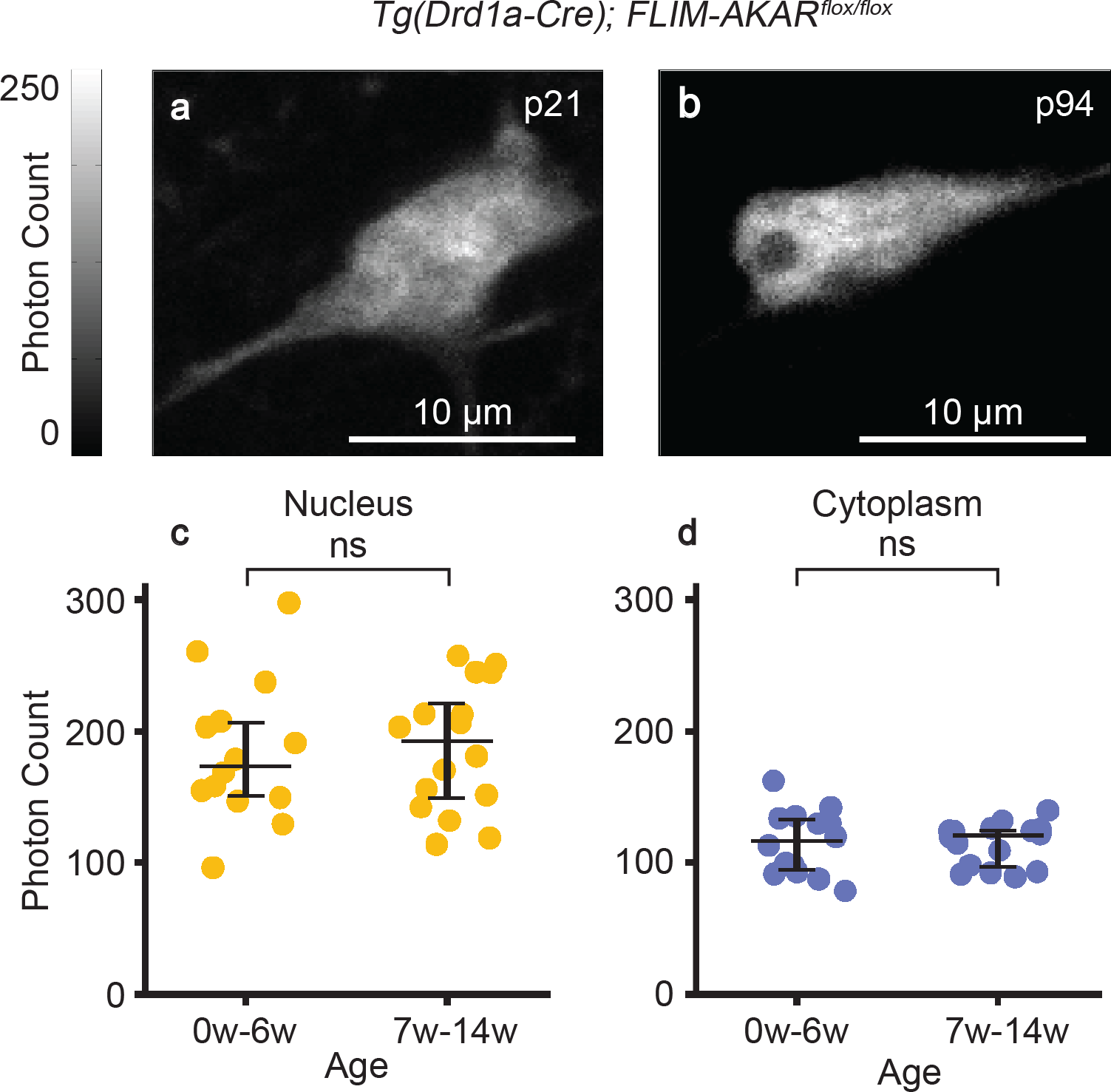
Stable expression of FLIM-AKAR in *FL-AK* reporter mouse over time. **(a-b)** 2p images of D1R-SPNs of the dorsal striatum in acute brain slices. (**c-d)** Quantification of nuclear and cytoplasmic photon count from two age ranges (0-6 weeks: n = 14 cells from 4 mice; 6-14 weeks: n = 16 cells from 6 mice). Data are represented as median with 25^th^ and 75^th^ percentiles (ns: not significant; p>0.05, 2-tailed Mann-Whitney U test).

### *FL-AK* mice report PKA activity in response to neuromodulator receptor activation in acute slices

PKA is activated by G_αs_-coupled receptors, one of the most well-known being the D1R. Signaling through D1Rs stimulates adenylate cyclase activity, which produces cyclic AMP (cAMP), a second messenger that can activate PKA^52,53^. D1Rs are found on various cell types, and play major functional roles in spiny projection neurons (SPNs) of the striatum^11,18,54–56^. SKF 81297 is a selective D1/D5 agonist that increases signaling through the cAMP/PKA pathway^54,56^. In order to assess whether *FL-AK* reporter mice can respond to D1 activation, we imaged FLIM-AKAR in D1R-SPNs of acute striatal slices from *Tg(Drd1a-Cre);FLIM-AKAR*^*f/f*^ mice. D1R-SPNs showed lifetime decreases in response to the D1/D5 agonist SKF 81297 (1 *μ*M), first in the cytoplasm and then in the nucleus, which is consistent with D1R activation beginning in the plasma membrane (Fig. 3a-c; nucleus: p = 1.30e-8, cytoplasm: p = 2.27e-12, Wilcoxon signed rank test). These results indicate that *FL-AK* reporter mice can report PKA activity increase in response to activation of a classical G_αs_-coupled receptor.

**Figure 3.**
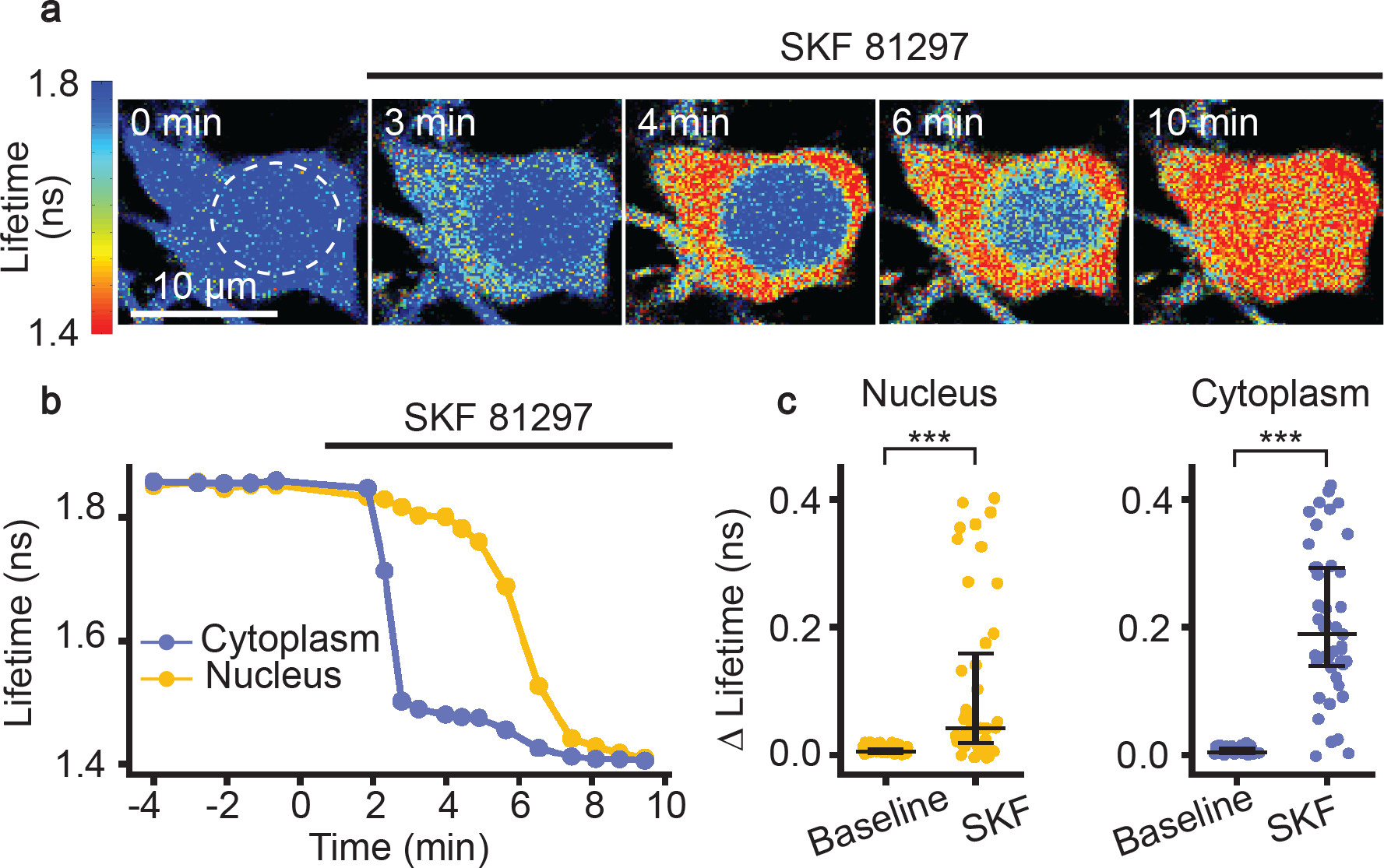
Functional response of D1R-SPNs to D1R activation in acute slices of *Tg(Drd1a-Cre); FLIM-AKAR*^*flox/flox*^ mice. **(a)** Time-lapse heatmaps of 2pFLIM images of an example SPN in acute slices of the dorsal striatum in response to the D1R agonist SKF 81297 (1 *μ*M). Dotted line indicates the location of nucleus. **(b)** Example trace of fluorescence lifetime response of the D1R-SPN shown in (a). **(c)** Quantification of change of fluorescence lifetime in response to SKF 81297 in D1R-SPNs (n = 43 cells from 6 mice). Data are represented as median with 25^th^ and 75^th^ percentiles (***: p <0.001, Wilcoxon signed rank test).

We subsequently determined whether *FL-AK* reporter mice can report elevated PKA phosphorylation in response to G_αq_-coupled receptor signaling. Although G_αq_ signaling was not classically linked to PKA, we recently discovered that endogenous G_αq_-coupled receptors, such as muscarinic acetylcholine receptors (mAChRs), do activate PKA^10^. Here, with *Emx1*^*IREScre*^*;FLIM-AKAR*^*flox/flox*^ mice, we assessed whether reporter mice expressing FLIM-AKAR in CA1 pyramidal neurons showed functional responses to muscarinic activation. Consistent with our previous data with IUE of the *FLIM-AKAR*^10^, we found a fluorescence lifetime decrease of FLIM-AKAR in both the nuclear and cytoplasmic compartments of CA1 pyramidal neurons in response to mAChR activation (Fig. 4a-c; nucleus: p = 0.0069, cytoplasm: p = 0.00066, Wilcoxon signed rank test; baseline vs muscarine). Following muscarinic receptor activation, we directly activated adenylate cyclase through the application of forskolin (FSK). In response, we saw an additional decrease in fluorescence lifetime, demonstrating a further increase in PKA activity (Fig. 4a-c; nucleus: p = 2.66e-6, cytoplasm: p = 9.31e-10, Wilcoxon signed rank test; muscarine vs muscarine + forskolin). These data indicate that *FL-AK* reporter mice can be used to detect changes in intracellular PKA activity in response to diverse neuromodulator inputs.

**Figure 4.**
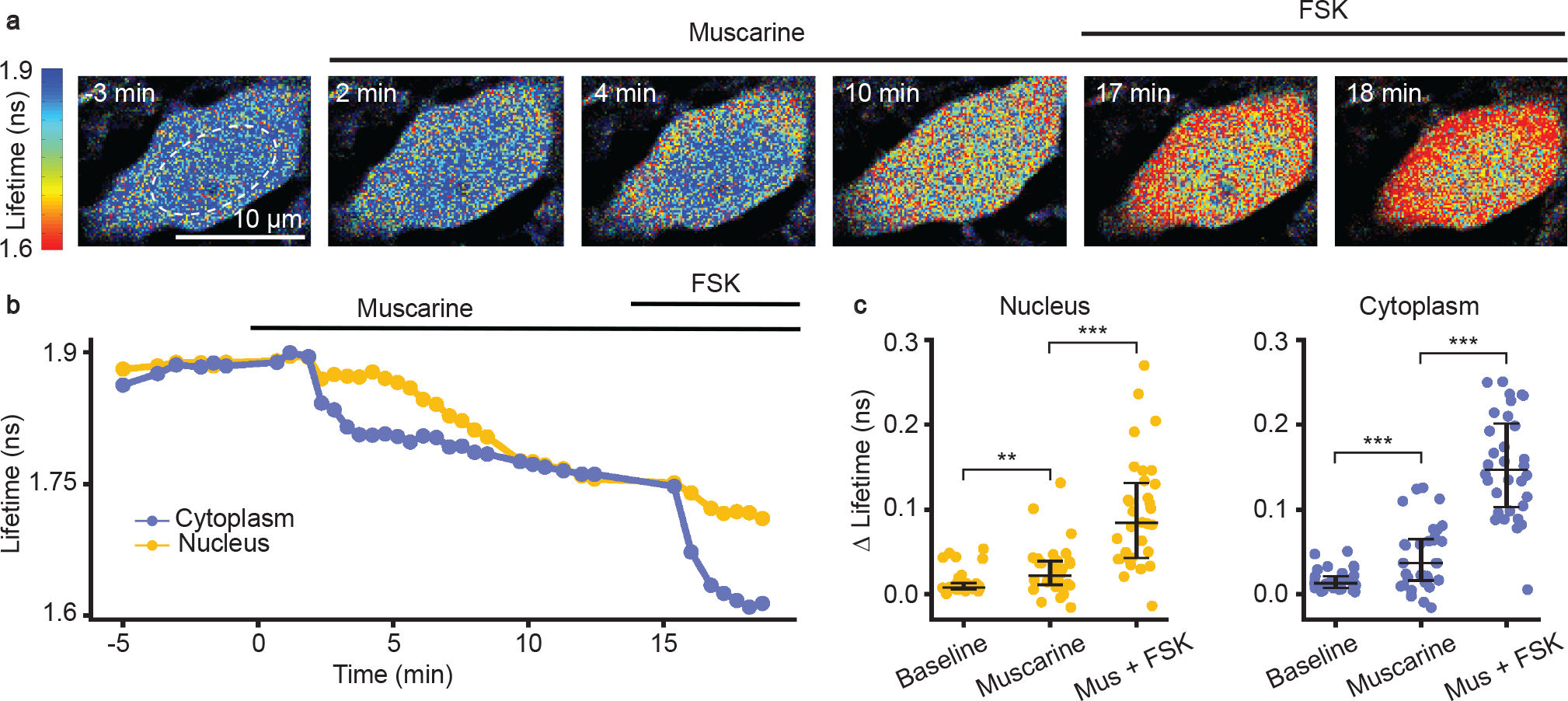
Functional response of CA1 pyramidal neurons in response to muscarinic acetylcholine receptor (mAChR) activation and adenylate cyclase activation in acute slices of *Emx1*^*IRESCre*^; *FLIM-AKAR*^*flox/flox*^ mice. **(a)** Time-lapse heatmaps of 2pFLIM images of an example CA1 pyramidal neuron in an acute hippocampal slice in response to the mAChR agonist muscarine (mus, 10 *μ*M) and adenylate cyclase activator forskolin (FSK, 50 *μ*M). Dotted line indicates the location of nucleus. **(b)** Example trace of fluorescence lifetime response of the pyramidal neuron shown in (a). **(c)** Quantification of change of fluorescence lifetime in response to mus and FSK in CA1 pyramidal neurons (n = 32 cells from 4 animals). Data are represented as median with 25^th^ and 75^th^ percentiles (***: p < 0.001, **: p<0.01, Wilcoxon signed rank test).

### *FL-AK* mice respond to dopamine receptor activation *in vivo*

We examined whether *FL-AK* reporter mice are sensitive enough to detect PKA activation *in vivo*. We implanted an optical fiber into the dorsal striatum of *Tg(Drd1a-cre);FLIM-AKAR*^*flox/flox*^ mice, and monitored PKA activity in freely moving mice with a custom fluorescence lifetime photometry (FLiP) setup^57^ (Fig. 5a). In response to intraperitoneal (IP) injection of the D1/D5 agonist SKF81297, but not saline, FLIM-AKAR in D1R-SPNs of the reporter mice showed a fluorescence lifetime decrease (Fig. 5b). These are consistent with previous reports where we delivered FLIM-AKAR with adeno-associated virus^57^. These results indicate *FL-AK* reporter mice can report PKA activity *in vivo*.

**Figure 5.**
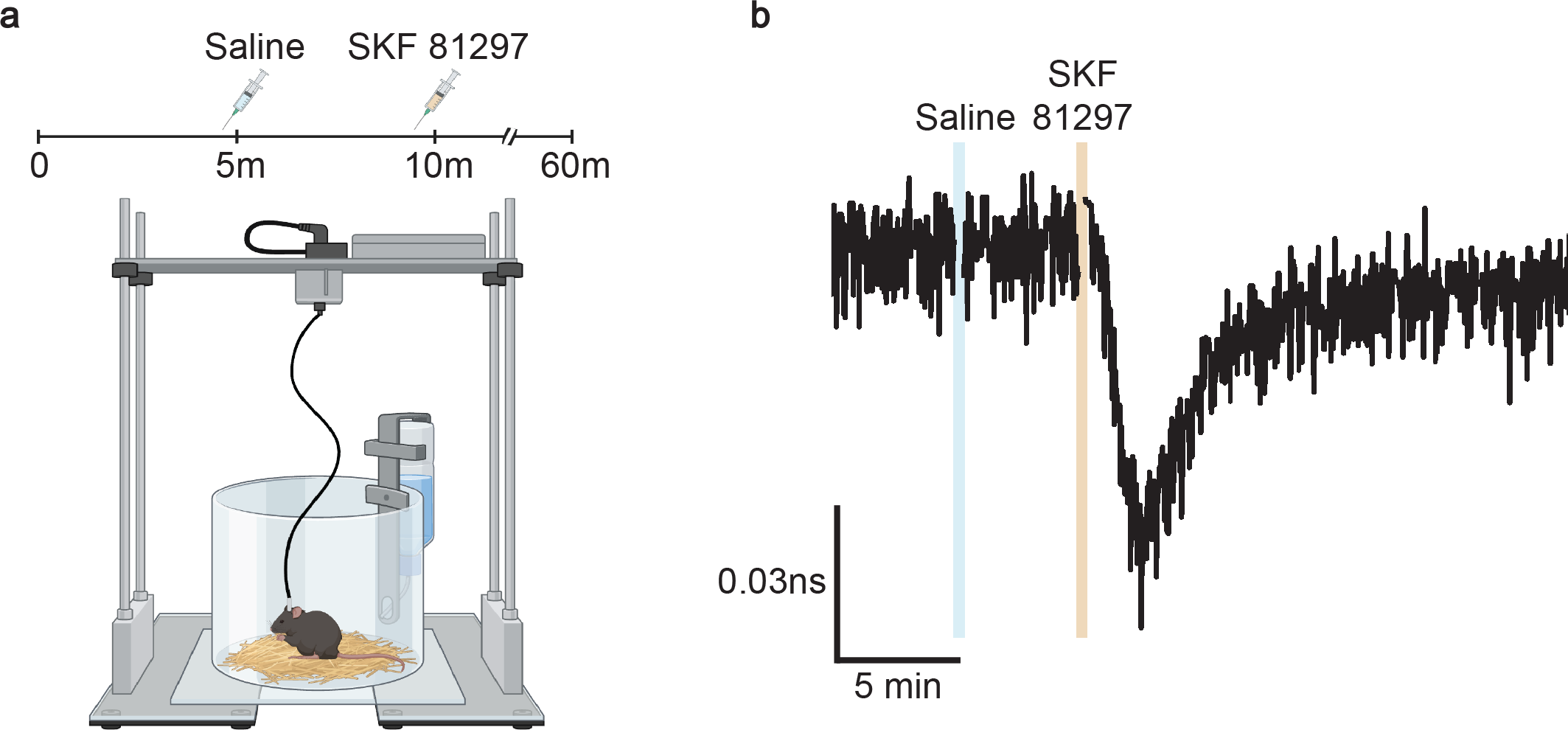
*In vivo* functional response of D1R-SPNs to D1R activation in *Tg(Drd1a-Cre); FLIM-AKAR*^*flox/flox*^ mice. **(a)** Schematic of experimental setup for fluorescence lifetime photometry (FLiP). **(b)** Example trace of change in fluorescence lifetime in response to saline followed by SKF 81297 (10 mg/kg) in D1R-SPNs in the dorsal striatum.

## Discussion

The FLIM-AKAR PKA activity reporter is a powerful optical tool that has the potential to unlock our understanding of the intracellular dynamics of PKA. *FL-AK* reporter mice facilitate the use of FLIM-AKAR in genetically defined cell populations specified by the *Cre* line with which the mice are crossed. Here, we show that *FL-AK*^*flox/flox*^ mice exhibit robust *Cre*-dependent expression of FLIM-AKAR that is stable over time. They also show reliable functional responses to diverse neuromodulator signals including activation of both G_αs_- and G_αq_-coupled receptors. Furthermore, *FL-AK* mice demonstrate a sufficient dynamic range to distinguish between different PKA phosphorylation states *in vivo*.

*FL-AK* mice offer several advantages over surgical methods to deliver the reporter. First, the *FL-AK* mouse line generates robust and consistent expression over time. Although fluorescence lifetime is largely insensitive to sensor expression levels, the variable expression seen in surgical delivery methods can result in differential contribution of autofluorescence, leading to an apparent sensor expression-dependent lifetime response. Thus, *FL-AK* reporter mice facilitate chronic imaging for transient PKA activation, comparison of basal PKA phosphorylation over time, and comparison of pooled results across multiple cells, mice, and experiments. Second, the mouse line eliminates the need for invasive surgeries like IUE or intracranial viral injection.

Third, although not explored in this study, *FL-AK* mouse line can allow effective targeting of sparsely distributed or hard-to-transfect cell types such as specific types of microglia and satellite glia. Fourth, this mouse line simplifies multiplex imaging to study how PKA activity changes in relation to other critical signaling molecules in cellular processes^58^.

Importantly, whereas this study focused on neuronal applications, *FL-AK* reporter mice have the potential to facilitate understanding the important roles of PKA dynamics in diverse tissues and body systems. For example, PKA has been studied in the context of immune modulation, cancer biology, and metabolic disorders such as obesity^27,28,33,59–64^. The delivery of *FLIM-AKAR* via surgical methods can pose a significant technical barrier to studying PKA dynamics in these tissues. Crossing *FL-AK* with diverse *Cre* lines will create opportunities to study PKA dynamics throughout the body and better understand how this signal modulates many critical processes.

## Materials and methods

### Knock-in mice

The floxed *FLIM-AKAR* reporter mouse line was generated by the Gene Targeting & Transgenics Facility at Howard Hughes Medical Institute’s Janelia Research Campus. *FLIM-AKAR* was knocked into the *ROSA26* locus, which was demonstrated to produce robust expression of inserted transgenes^65,66^. *CAG* promoter and the woodchuck hepatitis virus posttranscriptional regulatory element (WPRE) were used for robust and ubiquitous expression. A *lox-stop-lox* (LSL) cassette was included between the *CAG* promoter and the *FLIM-AKAR* open reading frame to produce *Cre* dependence. The *FLIM-AKAR* insert was cloned into a *ROSA26*-pCAG-loxp-STOP-PGKNeo-loxp-WPRE targeting vector^67^ between the second *loxp* and the *WPRE* sequences, where PGKNeo stands for polyphoglycerate kinase I promoter driving the neomycin phosphotransferase gene (PGK-Neo).

Aggregation method, where 8-10 embryonic stem cells were co-cultured with an 8-cell CD1 embryo, was used to produce chimeric mice.

Chimeric males were bred with CD1 female mice, and the female pups were crossed with the chimeric father to achieve a large number of pups for the homozygosity test to check for correct targeting. Genotyping of the floxed-FLIM-AKAR mice was performed by polymerase chain reaction (PCR) using the following primers:

R26 wt gt Forward: CCAAAGTCGCTCTGAGTTGT

R26 wt gt Rorward: CCAGGTTAGCCTTTAAGCCT

CMV scr Reverse: CGGGCCATTTACCGTAAGTT

PCR amplifies a fragment of 250bp of the endogenous *Rosa26* locus in wild type mice, and a fragment of 329bp between the *Rosa26* locus and the insert for floxed FLIM-AKAR mice.

The sperms of a correctly targeted F1 males were harvested, and *in vitro* fertilization of C57BL/6 females was performed by the Washington University Mouse Genetics Core. The progeny was bred with C57BL/6 mice for multiple generations to achieve strain stability and bred to homozygosity. The mice used in this manuscript were produced after 3 to 10 generations of breeding in a C57BL/6 background.

### Animals

All aspects of mouse husbandry and surgery were performed following protocols approved by Washington University Institutional Animal Care and Use Committee and in accordance with National Institutes of Health guidelines. The experiments were performed according to the ARRIVE guidelines^68^. *FLIM-AKAR*^*flox/flox*^ mice were crossed with *Emx1*^*IRESCre*^ (Jax: 005628)^51^ or *Tg(Drd1a-Cre)* (EY217Gsat; MGI: 4366803)^48–50^. For experiments examining expression over time, *Emx1*^*IRESCre*^; *FLIM-AKAR*^*flox/flox*^ mice were used (10 total mice: 8 females, 2 males; aged p21-p94). For experiments testing functional responses in acute brain slices, *Emx1*^*IRESCre*^; *FLIM-AKAR*^*flox/flox*^ were used to observe the response to muscarine (mus) in the hippocampus (4 total mice: 1 male, 3 females; aged p16-p19), and *Tg(Drd1a-Cre); FLIM-AKAR*^*flox/flox*^ mice were used to observe the response to SKF 81297 in the dorsal striatum (6 total mice: 3 males, 3 females, aged p33-p43). For *in vivo* studies, *Tg(Drd1-Cre); FLIM-AKAR*^*flox/flox*^ mice were used (1 mouse, female, aged 42 weeks).

### Implantation of Optical fibers

For FLiP experiments, an optical fiber (Doric Lenses, MFC_200/245-0.37_4.5mm_MF1.25_FLT) was implanted as described previously^69^. Here, the dorsal striatum was targeted using stereotaxic coordinates of 1.1 mm anterior and 1.7 mm lateral from Bregma and 2.5 mm ventral from the pia. Four stainless steel screws were secured in the skull to better anchor the dental cement.

### Acute Brain Slice Preparation

Acute brain slices were prepared as described previously^39^. For experiments involving *Tg(Drd1-Cre); FLIM-AKAR*^*flox/flox*^ mice, intracardial perfusion was performed with ACSF (final concentrations in mM: 127 NaCl, 25 NaHCO_3_, 1.25 NaH_2_PO_4_, 2.5 KCl, 1 MgCl_2_, 25 glucose) and slicing was performed in a cold choline-based cutting solution (final concentrations in mM: 25 NaHCO_3_, 1.25 NaH_2_PO_4_, 2.5 KCl, 7 MgCl_2_, 25 glucose, 0.5 CaCl_2_, 110 choline chloride, 11.6 ascorbic acid, 3.1 pyruvic acid) before slices were allowed to recover in ACSF at 34°C for 10 minutes. For experiments involving *Emx1*^*IRES-Cre/IRES-Cre*^*;FLIM-AKAR*^*flox/flox*^ mice, no intracardial perfusion was performed. Slicing was done in a cold sucrose-based solution (final concentrations in mM: 87 NaCl, 25 NaHCO_3_, 1.25 NaH_2_PO_4_, 2.5KCl, 75 sucrose, 25 glucose, 7.5 MgCl2) before the cells were allowed to recover in ACSF at 34°C for 10 minutes.

### Two-Photon Fluorescence Lifetime Imaging Microscopy (2pFLIM) and Image Analysis

2pFLIM was performed as previously described^69^ except for the following. 920 nm excitation wavelength was used to image FLIM-AKAR. For the calculation of lifetime, first, a double exponential curve was fitted to the lifetime histogram for the entire fields-of-view,

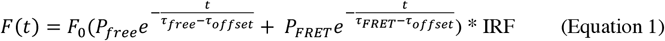

where *F(t)* is the photon count at lifetime t, *F*_0_ is the peak photon count, τ_*free*_ and τ_*FRET*_ are fluorescence lifetimes of donors that are free and that have undergone FRET respectively and are 2.14 ns and 0.69 ns for FLIM-AKAR. *P*_*free*_ and *P*_*FRET*_ are the corresponding fractions of these two species, τ_offset_ is the offset arrival time, and IRF is the measured instrument response function. Then, average lifetime for a given region of interest (ROI) was calculated from 0.0489 ns to 11.5 ns with the following calculation:

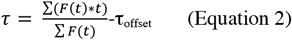

Where F(t) is the photon count at a given time channel, and t is the lifetime measurement at that time channel. Intensity measurement was represented by the photon count/pixel of a given ROI.

Change of fluorescence lifetime per cell at baseline was quantified as the absolute value of the difference between the average of the first three lifetime values of the baseline epoch and the minimum lifetime value of the baseline epoch. Change in lifetime due to drug treatment was quantified as the absolute value of the difference between the average of the last three lifetime values of the baseline epoch and the minimum lifetime value of the corresponding treatment epoch.

For Figure 2, all the data were collected at an imaging power of 2.5 mW and between 20-35*μ*m from the surface of the slice.

### Fluorescence Lifetime Photometry (FLiP) and Analysis

A FLiP setup was built and used as previously described^57,69^ except for the following. Fluorescence lifetime and intensity data were collected at 1Hz using our custom FLiP setup and acquisition software^57,69,70^. We calculated lifetime at each timepoint by first fitting a double exponential curve to the fluorescent lifetime histogram using a Gaussian IRF (Equation 1). Then, the average lifetime of the fitted curve (to infinity) was calculated.

Data were aligned to injection timepoints using synchronized video recordings through Bonsai (https://bonsai-rx.org/). Location of fiber implant was subsequently assessed with histology. Data analysis was performed using MATLAB.

### *In vivo* SKF81297 Response Experiments

Mice with an optical fiber implant were connected to a patch cord (Doric Lenses, MFP_200/220/900-0.37_1.5m_FCM-MF1.25_LAF) and placed in a round chamber to which they had been habituated previously. A camera was oriented to capture the entire recording chamber to assess behavior and capture injection time. Each trial consisted of 1 hour of continuous data collection with a saline injection (0.9% NaCl; 0.1 mL/10g body weight; IP delivery) at 5 minutes and an SKF 81297 injection (10 mg/kg) at 10 minutes.

Video recording was initiated in Bonsai at the start of FLiP recording using a transistor-transistor logic (TTL) pulse generated by Matlab through an Arduino Due board (Arduino, A000062) to ensure synchronized data collection. Video was collected at 25 frames per second.

### *In vitro* Pharmacology

For acute slicing experiments, drugs were applied via bath perfusion as previously described^69^. Final concentrations are indicated in parentheses: (+)-muscarine-iodide (10 *μ*M) and SKF 81297 hydrobromide (1*μ*M) were obtained from Tocris. FSK (50uM) was obtained from either Tocris or Cayman Chemicals.

### Quantification and Statistical Analyses

Detailed description of quantification and statistics are summarized in figure legends, figures, and results. Briefly, Mann-Whitney U test was used for unpaired data, Wilcoxon signed-rank test was used for paired data. Nonparametric tests were used so that we did not have to make any assumption of distribution. All tests were two-tailed with an alpha level of 0.05.

### Histology

For imaging of fixed whole-brain slices, both *Emx1*^*IRES-Cre/IRES-Cre*^*;FLIM-AKAR*^*flox/flox*^ and *Cre -/-;FLIM-AKAR*^*flox/flox*^ mice were used. Transcardiac perfusion was performed first with 1X Phosphate-Buffered Saline (PBS) and then with 4% paraformaldehyde (PFA) in PBS for tissue fixation^71^. Brains were placed in 4% PFA overnight before being switched to 1X PBS. 50*μ*m coronal sections were obtained with a vibratome (Leica Instruments, VT1000S). Sections were mounted on glass slides with mounting media containing DAPI stain. Images were obtained with a Zeiss Axioscan 7 using Zen Slidescan software. 475nm light was used for excitation. Images were acquired under both 5x and 10x objectives. Higher resolution images were subsequently stitched together.

## Data Availability

All data required to evaluate the conclusion are included in the Figures and text. All data will be freely available upon request for non-commercial purposes. The *FL-AK* mice will be available at the Jackson Laboratory Repository.

## Author Contributions

YC and BS conceived and designed the construction of the reporter mouse. ET, AM, and YC designed the characterization experiments. ET, AM, AO, and YC conducted experiments. ET, AM, and AO performed data analysis. ET and YC wrote the manuscript. All authors reviewed the manuscript.

## Additional Information

The authors declare no competing interests.

## Acknowledgments

We thank Caiying Guo, Mia Wallace, and members of Yao Chen’s lab for helpful feedback on the project. We thank Kerry Grens for critical comments on the manuscript. This work was supported by grants from the U.S. National Institute of Neurological Disorders and Stroke (R01 NS119821, to YC), the Whitehall Foundation (2019-08-64, to YC), and the U.S. National Institute of Aging (F30 AG084271, to ET). We thank Charles Gerfen and Nathaniel Heintz for the *Tg(Drd1a-Cre)* (EY217Gsat) mouse line. The floxed *FLIM-AKAR* reporter mouse line was generated by the Gene Targeting & Transgenics Facility at Howard Hughes Medical Institute’s Janelia Research Campus. In vitro fertilization was performed by the Mouse Genetics Core at Washington University School of Medicine. Data collection and analysis from experiments using the Zeiss Axioscan 7 and Zen Slidescan software were performed in part through the use of Washington University Center for Cellular Imaging (WUCCI) supported by Washington University School of Medicine, The Children’s Discovery Institute of Washington University and St. Louis Children’s Hospital (CDI-CORE-2015-505 and CDI-CORE-2019-813) and the Foundation for Barnes-Jewish Hospital (3770 and 4642). Schematic illustrations from Fig. 1a-b and 5a were created with BioRender.

## Notes

### Competing Interest Statement

The authors have declared no competing interest.

